# A conserved metabolic signature associated with response to fast-acting antimalarial agents

**DOI:** 10.1101/2022.10.03.510714

**Authors:** Nelson V. Simwela, W. Armand Guiguemde, Judith Straimer, Clement Regnault, Fumiaki Yokokawa, Benjamin Taft, Thierry T. Diagana, Michael P. Barrett, Andrew P. Waters

## Abstract

Characterizing the mode of action of antimalarial compounds that emerge from high-throughput phenotypic screens is central to understanding how parasite resistance to these drugs can emerge. Here, we have employed untargeted metabolomics to inform on the mechanism of action of antimalarial leads with different speed of kill profiles being developed by the Novartis Institute of Tropical Diseases (NITD). Time-resolved global changes in malaria parasite metabolite profiles upon drug treatment were quantified using liquid chromatography-based mass spectrometry (LC-MS) and compared to untreated controls. Using this approach, we confirmed previously reported metabolomics profiles of the fast-killing (2.5h) drug dihydroartemisinin (DHA) and the slower killing atovaquone (ATQ). A slow acting antimalarial lead from NITD of imidazolopiperazine (IZP) class, GNF179, elicited little or no discernable metabolic change in malaria parasites in the same 2.5h window of drug exposure. In contrast, fast killing drugs, DHA and the spiroindolone (NITD246) elicited similar metabolomic profiles both in terms of kinetics and content. DHA and NITD246 induced peptide losses consistent with disruption of haemoglobin catabolism and also interfered with the pyrimidine biosynthesis pathway. Two members of the recently described novel class of antimalarial agents of the 5-aryl-2-amino-imidazothiadiazole (ITD) class also exhibited a fast-acting profile that also featured peptide losses indicative of disrupted haemoglobin catabolism. Our screen demonstrates that structurally unrelated, fast acting antimalarial compounds generate similar biochemical signatures in *Plasmodium* pointing to a common mechanism associated with rapid parasite death. Our study describes a potential biochemical signature that may serve to identify other fast acting drug candidates.

**Importance:** In malaria drug discovery, understanding the mode of action of lead compounds is important as it helps in predicting the potential emergence of drug resistance in the field when these drugs are eventually deployed. In this study, we have employed metabolomics technologies to characterize the potential targets of antimalarial drug candidates in the developmental pipeline at NITD. We show that NITD fast acting leads belonging to spiroindolone and imidazothiadiazole class induce a common biochemical theme in drug exposed malaria parasites which is similar to another fast acting, clinically available drug, DHA. These biochemical features which are absent in a slower acting NITD lead (GNF17) point to haemoglobin digestion and inhibition of the pyrimidine pathway as potential action points for these drugs. These biochemical themes can be used to identify fast drug candidates of similar profiles in future drug discovery programs.

## Introduction

Malaria is a resurgent worldwide public health problem affecting millions and in the shape of *Plasmodium falciparum* killing hundreds of thousands of people annually (1). Central to malaria control programs are antimalarial drugs, which form crucial components of the current malaria treatment, prophylaxis and transmission blocking strategies. Artemisinins (ARTs), in artemisinin combinational therapies (ACTs), have been the backbone of these strategies in the last decade and have contributed significantly to the recent gains achieved in malaria control (1, 2). ACTs currently remain mostly effective in Sub-Saharan Africa, a region that harbours the highest burden of the disease. However, as has been the historical trend with antimalarial treatments, resistance to ARTs emerged in South East Asia (SEA) and was first reported in 2009 along the Thai-Cambodian border (3). At present, ART treatment failure has reached endemic status in SEA (1, 4, 5) and has seemingly emerged in Africa (6). More worryingly, parasites carrying both ART and partner drug piperaquine resistance mutations, have been reported in SEA threatening the current mainstay of ACTs (7, 8). Pipelines to identify new drugs to combat the emerging resistance or for effective combination therapies are thus urgently needed.

Over the past decade, thousands of chemical entities that block malaria parasite growth have been reported from pharmaceutical companies and public funded product development partnerships (9-11). These screens have provided appropriate starting points for antimalarial drug discovery which could serve as potential replacements and/or suitable combination partners with current drugs to combat and overcome resistance. However, as is the case with a majority of anti-parasite compounds identified through phenotypic screens against parasites, their mode of action is unknown (12, 13). Characterising the mode of action of lead drug candidates, or drugs which are already in clinical use, though not essential during drug development, is important as it provides a platform to understand or predict resistance mechanisms as well as identify suitable combination drug partners using mode of action informed strategies. Mode of action elucidation also helps in identifying the actual drug targets which can be exploited in structure-based design towards better drugs.

Mode of action elucidation in malaria parasites has primarily involved forward genetics approaches, which involve *in vitro* selection for resistance followed by whole genome sequencing, transcriptome and or proteome analysis. These approaches have identified or confirmed the mechanism of resistance and mode of action of known antimalarial drugs such as ATQ, pyrimethamine and chloroquine (14-16). They have also pointed to potential molecular targets of novel compounds e.g. phosphatidylinositol 4-kinase (17), protein synthesis (18) and the pyrimidine biosynthesis pathways (19). These forward genetic screens, however, have their own limitations as they might not reveal the full range of molecular and biochemical networks involved in drug resistance processes (13). Moreover, these screens cannot be used to probe mode of action of chemical entities when drug resistance cannot be selected (high barrier compounds), drug resistance is phenotypic (20) or when resistance is conferred by gene mutations in multi-drug resistance transporters which provide little or no clue as to the intracellular target of the compounds. Metabolomics screening platforms now provide an alternative approach to elucidating the mode of action of both known drugs and lead candidates in bacterial pathogens (21, 22) and malaria parasites (23-25) as many antimicrobial agents target metabolic enzymes and pathways. This has been made possible because metabolomics platforms can detect perturbations induced by drug treatment under controlled *in vitro* exposure conditions (13, 21, 26). In malaria parasites, these approaches have been used to identify the mode of action of polyamine inhibitors (27), to validate the activity of new quinolone drugs targeting the parasite electron transport chain (28) and to reveal metabolic specific phenotypes associated with the clinically relevant drugs dihydroartemisinin (DHA) and chloroquine (24, 29). Metabolomic screens of the malaria box compounds have also revealed established as well as novel targets of potential malarial drug candidates (23, 25). Combinatorial -omics approaches can provide even greater detail, as, for example, high-resolution metabolomics combined with peptidomics and biochemical analyses revealed that a novel fast-acting lead drug candidate being developed by the Medicines for Malaria Venture (JPC-3210, MMV 892646) possibly acts by inhibiting haemoglobin catabolism and protein translation (30).

In this study, an untargeted metabolomics approach was used to screen two, novel fast acting drug candidates of the ITD class that have emerged from the NITD drug discovery pipelines (31). We first validated our metabolic profiles using ATQ which has a well characterised metabolic fingerprint (23, 24). Thereafter the metabolic profile of two representatives of the ITD compound series; compound 9 (Cpd 9) and compound 55 (Cpd 55) (31) was compared to other drug candidates from the NITD pipeline; spiroindolones which are known to target the *P. falciparum* Na^+^ H^+^ ATPase (PfATP4) (32) and GNF179 (KAF156 analogue) whose precise mode of action is still unknown (33-35). These profiles were further compared to that of the lead fast acting antimalarial DHA using a fixed time point as well as a dynamic time course over the first 2.5 hours of drug exposure. We demonstrate that the metabolic profile of the fast-acting antimalarial drugs and drug candidates tested here, of greatly differing chemical structure reveal fundamental similarities in their impact on the parasite, interfering with haemoglobin catabolism. We conclude that this could be a common metabolic response of the parasites which reflects either a commonality in the mode of action or a common parasite response to rapidly induced death.

## Results and discussion

### ITDs display a fast-killing rate

Parasite killing rates allow for identification of fast acting compounds which are desirable for malaria control as they allow for rapid clearance of parasitaemia in patients, which in turn minimises parasite drug exposure time and narrows the window in which parasites can evolve resistance. *In vitro* assays to predict the parasite killing rates of antimalarial compounds are based on parasite reduction ratios (PRR), quantified over 28 days by fresh exposure of a defined parasite inoculum every 24 hours in a series of limiting dilutions (36). Even though the PRR method allows for determination of parasite clearance times as well as drug lag phases (the time required for compounds to achieve maximum killing effect), the extent to which parasite metabolic and biochemical fingerprints change, especially for potentially pleiotropic, fast acting compounds where the onset of action is less than 24 hours, cannot be predicted accurately. We determined the killing kinetics of two ITD series of compounds (Cpd 9 and Cpd 55), NITD246, DHA, GNF179 and ATQ by biochemically monitoring luciferase expression using a *P. falciparum* 3D7 luc line which constitutively expresses a dual NanoLuc and luciferase reporter. Synchronised trophozoites at 2% parasitaemia and 2% haematocrit were cultured with the compounds at 10 × 72 hours IC_50_ (**Table 1**) or DMSO (0.1%) after which the relative luminescence signal (RLU) was monitored over the course of 6 hours. DHA (1 uM), which is a known fast acting compound, depleted the luciferase signal after 2.5 hours of incubation (**Fig. 1A**). A member of the spiroindolone series (NITD246) also depleted the signal after 2.5 hours but at a faster initial rate than DHA (**Fig. 1A**). This is in agreement with observations that spiroindolones exert a faster mode of action and parasite clearance than ARTs in patients (37). Meanwhile, both Cpd 9 and Cpd 55 at 10 x IC_50_ displayed an even faster killing rate, depleting the luciferase signal after 2 hours of drug incubation in line with previous observations in the PRR assays (31). By contrast, GNF179 and ATQ which are known to act slowly had a negligible effect on the luciferase signal over 6 hours at 10 x IC_50_, displaying an almost identical response to the DMSO control. Microscopic analysis of parasite morphologies during the time points did not show any significant differences relative to the DMSO control (**Fig. 1B**). Based on these data, we chose 2.5 hours as our initial first time point for metabolomics drug incubation as it was the time which corresponded with maximal biochemical signal disruption (based on luciferase expression) for the fast-acting compounds and has been previously shown to yield a good metabolic signal even for slow acting compounds such as ATQ (23).

**Table 1.**
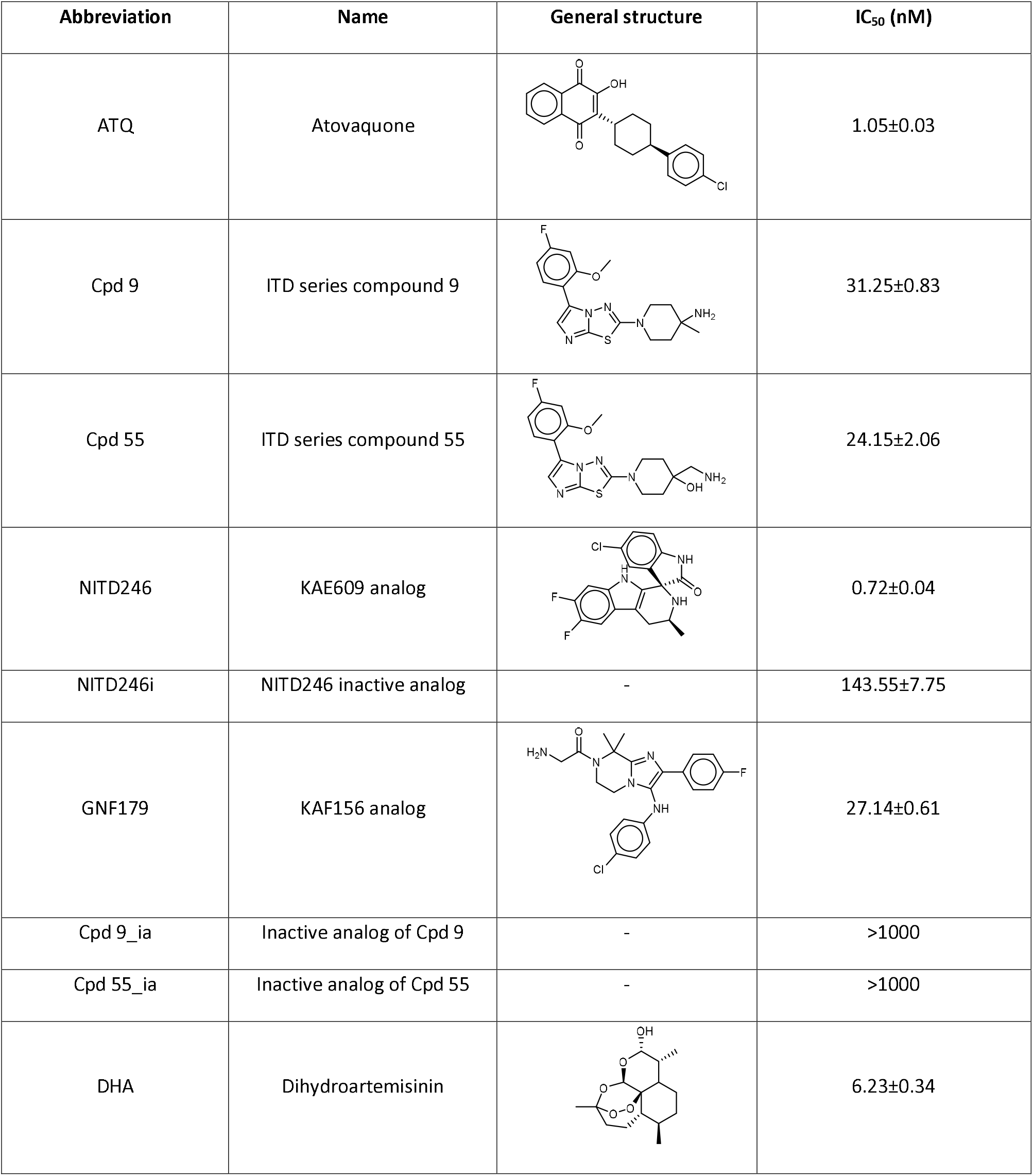
List of antimalarial compounds used in this study, their structures and IC_50_ values. IC_50_ values are means and standard deviations from three biological repeats.

| Abbreviation | Name | General structure | IC<sub>50</sub> (nM) |
| --- | --- | --- | --- |
| ATQ | Atovaquone |  | 1.05±0.03 |
| Cpd 9 | ITD series compound 9 |  | 31.25±0.83 |
| Cpd 55 | ITD series compound 55 |  | 24.15±2.06 |
| NITD246 | KAE609 analog |  | 0.72±0.04 |
| NITD246i | NITD246 inactive analog | - | 143.55±7.75 |
| GNF179 | KAF156 analog |  | 27.14±0.61 |
| Cpd 9_ia | Inactive analog of Cpd 9 | - | >1000 |
| Cpd 55_ia | Inactive analog of Cpd 55 | - | >1000 |
| DHA | Dihydroartemisinin |  | 6.23±0.34 |

**Fig. 1:**
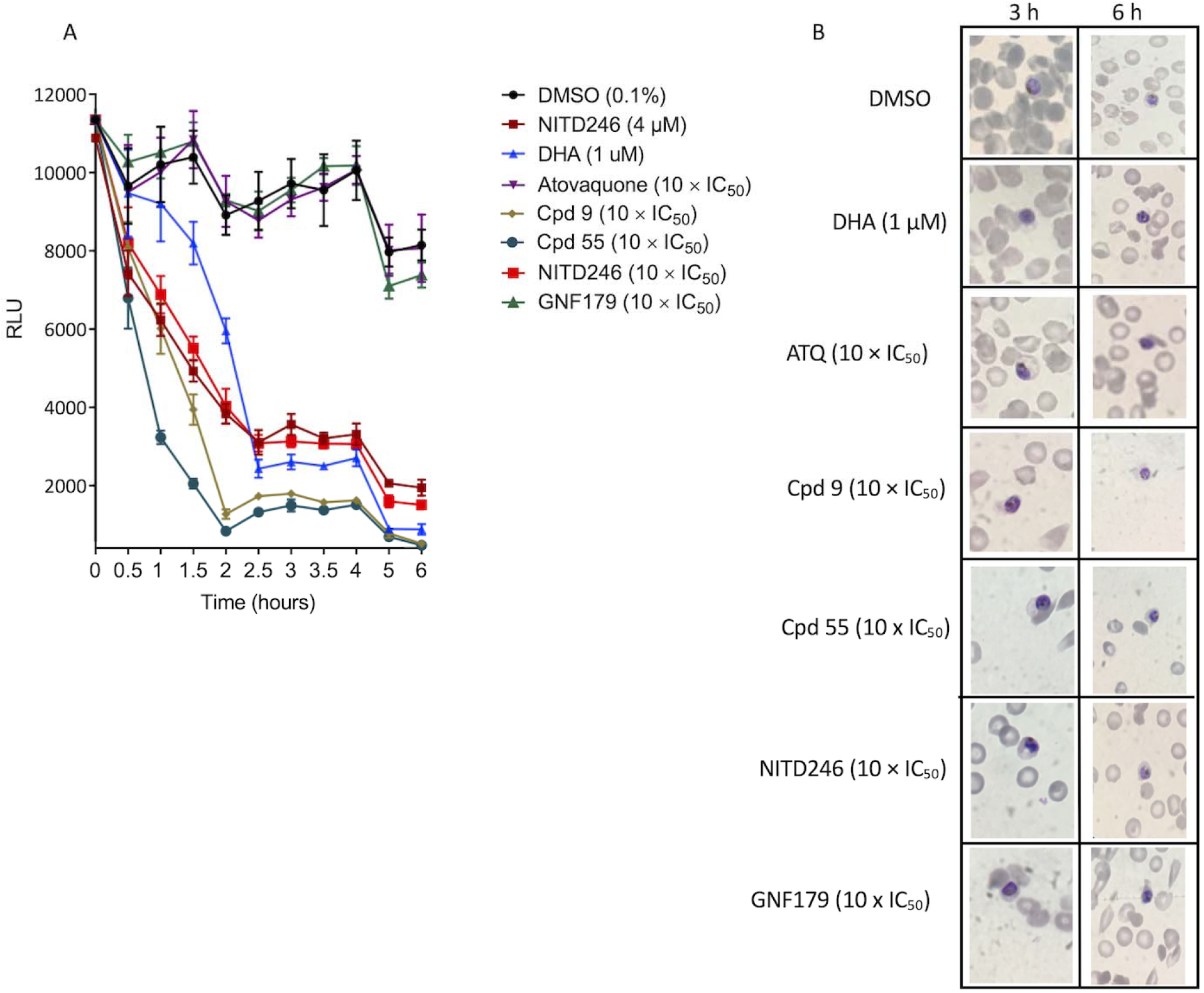
Killing kinetics of NITD246, DHA, ATQ, Cpd 9, Cpd 55 and GNF179 in the 3D7 luc line. ∼30-hour old trophozoites at 2% haematocrit and 2% parasitaemia were incubated with the compounds at the indicated concentrations for the indicated times. Luciferase expression was quantified at each time point. **A**. plot of relative luminescence unit (RLU) over the 6-hour incubation periods for the compounds. Incubations were carried out in quadruplicate over two independent biological repeats. **B**. Microscopy analysis of Giemsa-stained smears at the 3- and 6-hour incubation periods for all the compounds.

### ATQ disrupts pyrimidine biosynthesis pathway in malaria parasites

ATQ targets the mitochondrial electron transport chain (mETC) bc1 complex (Cbc1) that plays a crucial role in oxidative phosphorylation in most organisms (38). However, malaria parasites do not require oxidative phosphorylation and have maintained an active mETC in the asexual blood stages for the sole purpose of recycling ubiquinone, which acts as an electron acceptor for dihydroorotate dehydrogenase (DHODH), a critical enzyme in the pyrimidine biosynthesis pathway. The parasite’s mETC also indirectly feeds into the Tricarboxylic Acid pathway (TCA) by supplying ubiquinone which is essential for the activity of dehydrogenases of the TCA, succinate dehydrogenase (SDH) and malate dehydrogenase (MDH) (39). Purified trophozoites exposed to ATQ at 10 x IC_50_ for 2.5 hours, revealed a rapid accumulation of N-carbamoyl L-aspartate and dihydroorotate while the level of downstream pyrimidine metabolites; uridine diphosphate (UDP) and uridine triphosphate (UTP) declined (**Fig. 2A, 2B, Fig. S1B, S1C**). This is in agreement with previously reported metabolomic profiles for ATQ (23, 24). We also observed that orotate levels were maintained despite DHODH inhibition with ATQ (**Fig. S1A**) as has also been previously observed (24). The mechanism underlying this phenomenon remains unclear. ATQ treatment also led to a decrease in cellular levels of citrate (**Fig. 2B**). Blood stage malaria parasites appear to carry out a canonical oxidative TCA where the majority of the carbon enters the cycle as α-ketoglutarate derived from a series of glutaminolytic reactions (40). However, a low flux of glucose derived carbon into the TCA through acetyl-CoA has also been reported (40, 41). Meanwhile, it has been demonstrated that ATQ resistance (a bypass of mETC inhibition) can be achieved in transgenic malaria parasites that artificially express yeast DHODH which does not require ubiquinone for its activity (39). Conversely, mETC inhibition by ATQ acts as a de facto knockout of all enzymes that require ubiquinone for activity which include DHODH, SDH and MDH. Indeed, using the same parasites expressing yeast DHODH which can tolerate otherwise lethal doses of ATQ, stable isotope labelling was used to demonstrate that ATQ activity does not just inhibit the mETC, but also prevents flux of glucose derived carbons into the TCA (42). Previous ATQ metabolomics profiles have also revealed disruption in the TCA cycle as an accumulation of fumarate was observed, consistent with interference of SDH or MDH, as a consequence of Cbc1 complex inhibition (24). Even though we do not observe an accumulation of fumarate in our metabolomics screen (**Fig. S1D**) which could be due to shorter drug exposure time, a steady decrease in the TCA cycle metabolites, which was previously comprehensively profiled upon stable isotope labelling (24), is mirrored by the observed decrease in levels of citrate in our profiling. Taken together, these results validated our metabolomics approach for mode of action elucidation of selected NITD drug candidates (**Table 1**).

**Fig. 2:**
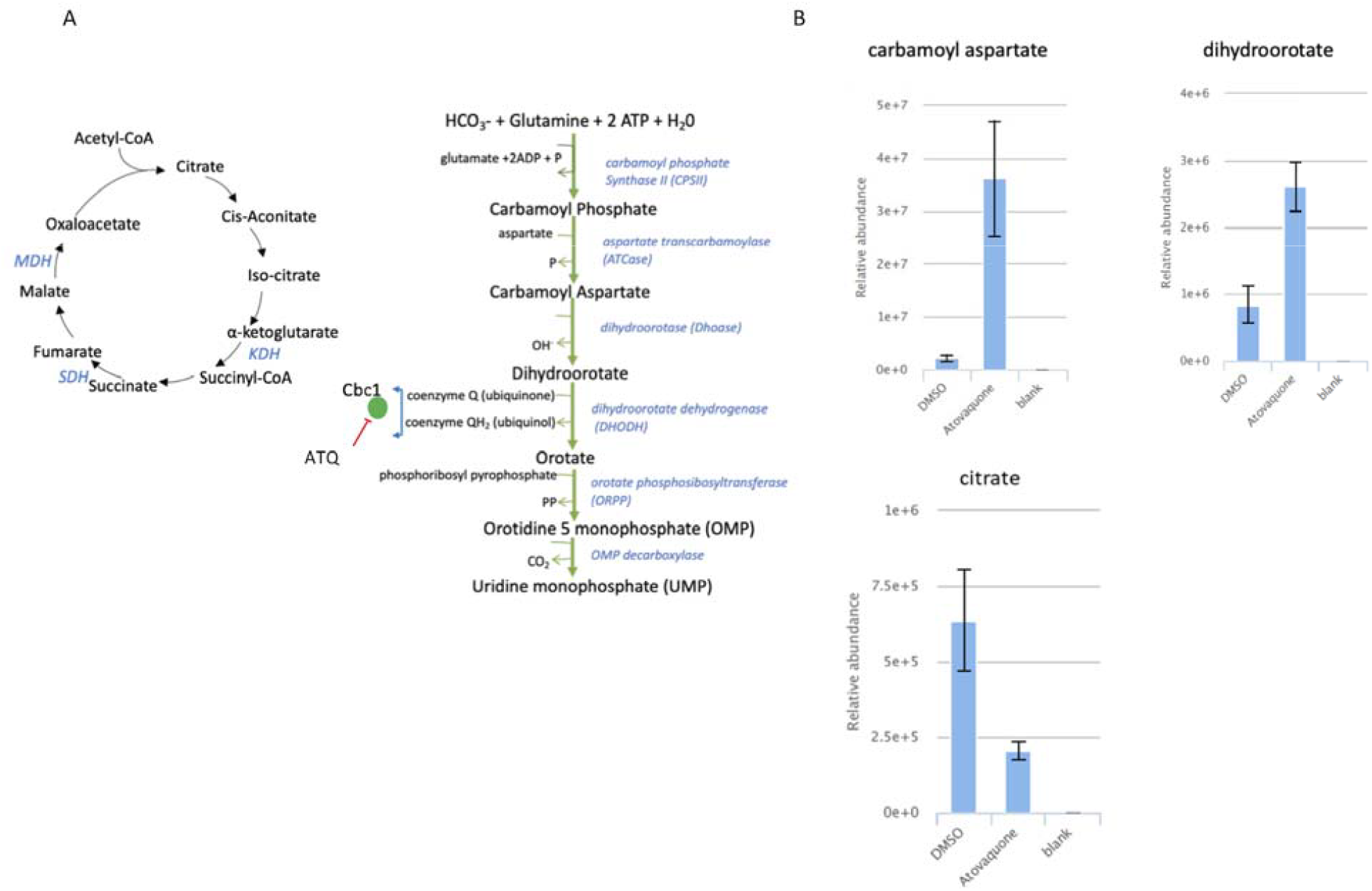
Metabolomics profile of ATQ. ∼1 × 10 purified trophozoites were exposed to either DMSO or ATQ at 10 X IC_50_ for 2.5 hours. Untargeted metabolomics on an LC-MS platform was carried out on extracted metabolites. **A**. Schematic of the TCA cycle and pyrimidine pathways. The cytochrome bc1 complex (Cbc1) which is the target of ATQ is shown in relation to its role in recycling of ubiquinol for DHODH activity and regeneration in the TCA by indicated dehydrogenases. KDH-ketoglutarate dehydrogenase, SDH-succinate dehydrogenase, MDH-malate dehydrogenase (also called malate quinone oxidoreductase). **B**. Relative abundance of the indicated pyrimidine and TCA metabolites in DMSO vs ATQ treatments. Relative abundance measurements are comparisons of total ion counts of the metabolites in the treatment conditions. Treatments were carried out in triplicates over two independent biological repeats. mzXML mass spectrometry files and graphs were processed and plotted in pIMP

### NITD246 elicits a pleiotropic metabolic response in malaria parasites

NITD246, a spiroindolone analog of KAE609, is one of the fast-acting compounds developed by NITD and has shown promising results in clinical trials (37, 43). Using forward genetic screens after *in vitro* selection for resistance, spiroindolone KAE609 was proposed to target PfATP4, a Na^+^ H^+^ ATPase, even though the exact events preceding parasite death remain unknown (32). Treating malaria parasites with KAE609 has been shown to lead to a rapid influx of sodium, increased rigidity of infected red blood cell membranes and consequent alteration of parasite morphology/rheology (44, 45). We compared the metabolomic profile of *P. falciparum* exposed to NITD246 or its inactive analogue NITD246i after 2.5 hours of incubation with 10 x IC_50_ of NITD246. A massive reduction in peptide levels (many of them potentially haemoglobin derived) was observed (**Fig. 3A, Table S1**). Moreover, NITD246 incubation resulted in accumulation of choline and glycero-phosphocholine (**Fig. 3B**), disrupted the pyrimidine biosynthesis pathway (but with a different signature to ATQ, **Fig. 3C, 3D, Fig. S2A**) and also caused a loss in purine metabolites (**Fig. S2B**). These data are in contrast to the inactive analogues which yielded profiles similar to the DMSO control. These observations are also similar to previous metabolomic profiles for KAE609 which reported a loss of haemoglobin derived peptides, amino acid derivatives and central carbon metabolites (23). This illustrates a potential pleiotropic metabolic response which could arise as a result of rapid disruption of cellular homeostasis upon PfATP4 inhibition and sodium influx. In spite of the evidence indicating that KAE609 targets PfATP4, it remains questionable whether PfATP4 is the direct target of this compound or acts as a multidrug resistance gene. Mutations in PfATP4 do not just confer resistance to KAE609, but to a diverse array of chemically unrelated compounds (such as carboxymides, aminopyrazoles, pyrazoleamides and dihydrosioquinolines) which all possess potent antimalarial activity (46-48). Our metabolomics data point to primary or secondary pleiotropic events associated with exposure to KAE609, including diminished haemoglobin catabolism and the inhibition of pyrimidine biosynthetic pathways which could be part of the mode of action or consequence of NITD246 mediated parasite killing. Interestingly, although the pyrimidine metabolic fingerprint of NITD246 differs from that of ATQ (**Fig. 3D**), it is similar to the previously reported pyrimidine metabolomic profile of DHA treated parasites, albeit at differing time points (24). These data suggest that both DHA and spiroindolones directly or indirectly perturb the early enzymes of pyrimidine biosynthesis (ATCase, CPSII) leading to diminished levels of downstream metabolites. In a separate metabolomic profiling of malaria box compounds (25), metabolic fingerprints of DHA treated parasites were shown to cluster together with KAE609 and also some other PfATP4 inhibitors; SJ733 and MMV006427. The actual events leading to parasite death in NITD246 could thus potentially involve promiscuous targeting, similar to DHA, shutting down multiple biological pathways in the parasite.

**Fig. 3:**
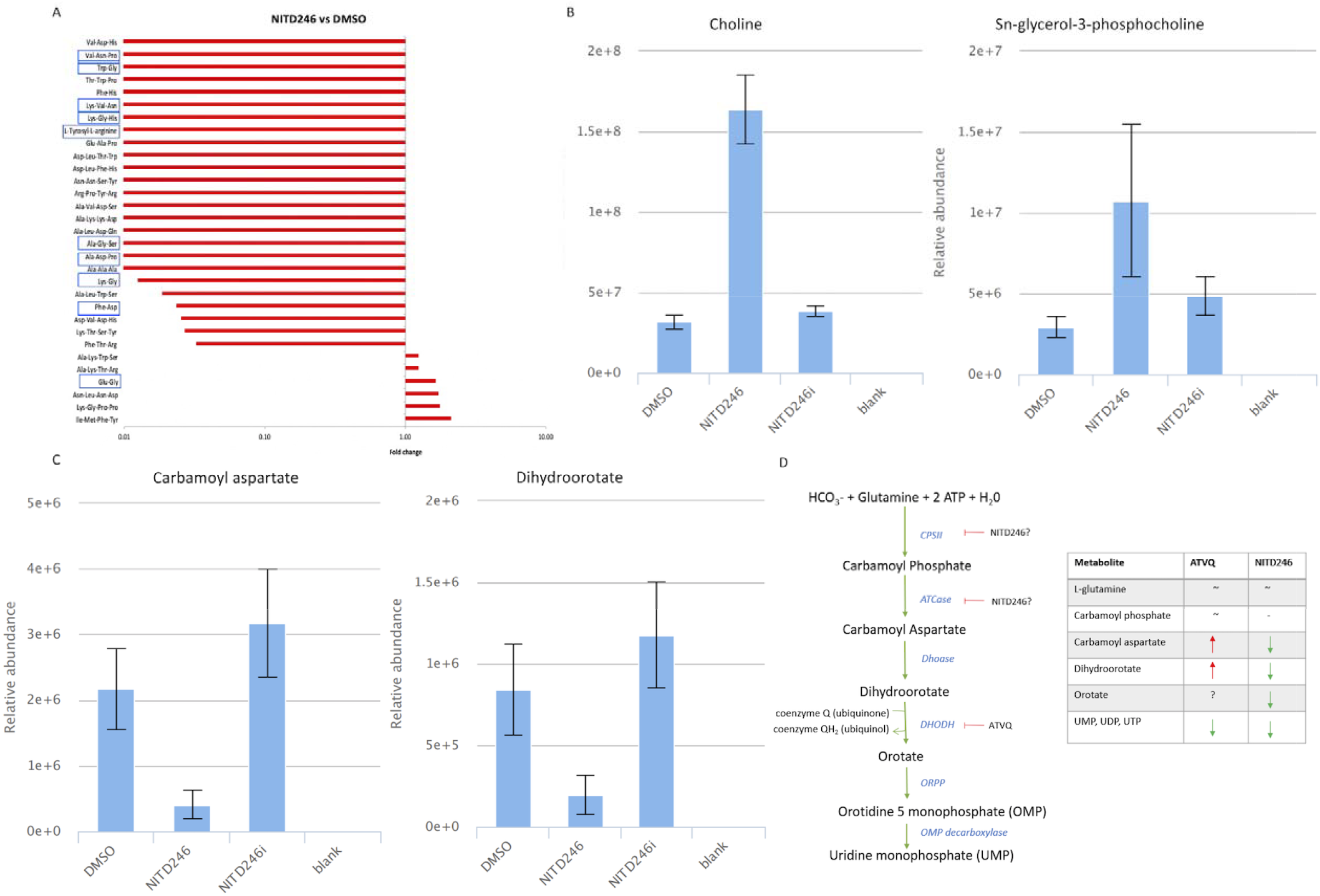
Pleiotropic metabolic response of malaria parasites after exposure to NITD246 for 2.5 hours. **A**. Global untargeted metabolomic response of selected peptides in NITD246 treated trophozoites as compared to DMSO. Peptides with haemoglobin matching sequences as well as those which are potentially haemoglobin derived (23-25) are highlighted with blue rectangular shapes. **B**. Relative abundance of choline and choline derivatives in the NITD246 treated parasites as compared to DMSO and in-active analogue. C. Relative abundance of the indicated pyrimidine biosynthesis pathway metabolites in the NITD246 treated parasites as compared to DMSO and in-active analogue. D. Cartoon of the pyrimidine biosynthesis pathway showing ATQ (Fig. 2) and NITD246 potential action points. The table is a direct comparison of ATQ and NITD246 pyrimidine biosynthesis pathway metabolites. Fold changes (relative to DMSO control) and relative abundance comparisons are means from two biological repeats collected in triplicate at each time of drug incubation.

### ITD series compounds elicit minimal metabolic responses

Cpd 9 and Cpd 55 are analogues of the 5-aryl-2-amino-imidazothiadiazole class of compounds that are in the developmental pipeline at the NITD (31). They are very fast acting (Cpd 9 t½ ∼1h; Cpd 55 t½ < 45min), even more rapid than spiroindolones (NITD246) and DHA (**Fig. 1A**), (31). Attempts to generate parasite lines highly resistant to these compounds by *in vitro* selection have so far been unsuccessful, making attempts to characterise their mode of action particularly difficult. In our metabolomics screen, Cpd 9 and Cpd 55 induced very similar metabolic profiles, which were almost entirely restricted to significant reductions across predicted peptides (**Fig. 4A, 4B, Table S2, S3**). Some of these peptides (highlighted in **Fig. 4A, 4B**) could be mapped to the α and β chain sequences of haemoglobin, a similar perturbation to that reported previously in metabolic profiling of DHA (24, 25). This would suggest that ITDs potentially target haemoglobin breakdown as they exert their anti-parasite activity. The ITD peptide response appears to be similar to those observed with NITD246 as well as DHA in previous screens (23-25). While DHA has been proposed to target haemoglobin catabolism, NITD246 is believed to target PfATP4 (32) hence the haemoglobin breakdown response associated with exposure to this drug may be secondary. The ITD peptide response could also be a secondary consequence of related inhibition of a target not revealed by our metabolomics screen. Moreover, the true source of short di/tripeptides revealed here are difficult to ascertain with certainty. For example, the Met-Ala, Trp-Pro, Leu-Met peptide combinations which are also significantly perturbed in the ITDs metabolic profile (**Fig. 4A, 4B**), are not present in haemoglobin sequences. This could point towards a more general inhibition of protein degradation systems, impaired flux of oligopeptides due to inhibition of transporters (49) or a signal of dying parasites that bears no relevance to the target or primary mode of drug action. Several compounds with unrelated primary modes of action (DHA, spiroindolones, aminopyridines; PI4K inhibitors, triaminopyrimidines; vacuolar ATPase inhibitors, chloroquine) have been shown to perturb haemoglobin catabolism (23, 50). However, the ITD peptide response we note here is not accompanied by disruption to other pathways such as pyrimidine or purine nucleotide responses, as is the case with NITD246 or DHA. Interestingly, another fast-acting compound which has been developed by the MMV, JPC-3210, elicits a similar metabolic response to the ITDs, i.e. being mostly restricted to peptides leading to the proposal that inhibited haemoglobin catabolism is its primary mode of action, although interference with protein translation was also noted which might as well be a feature of ITD activity (30).

**Fig. 4.**
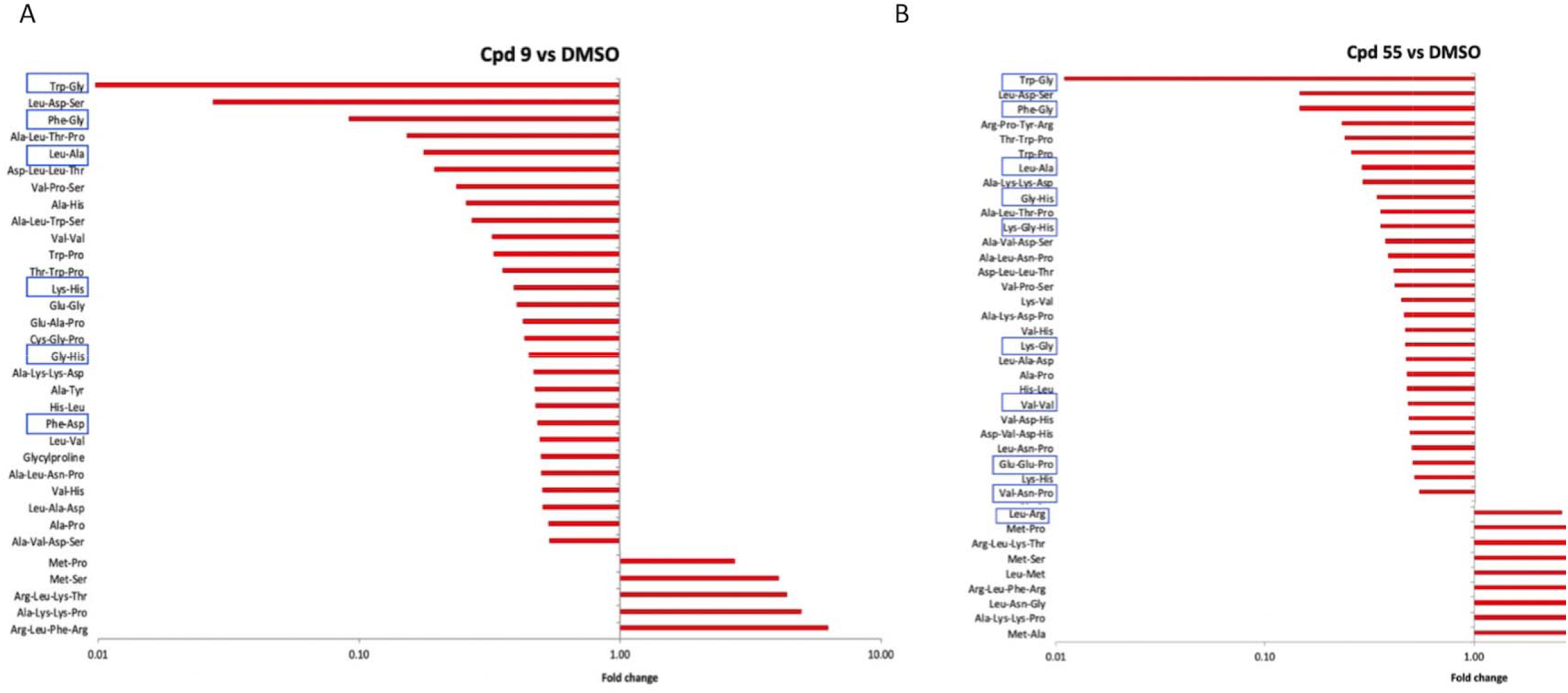
Peptide metabolic responses of the ITD series compounds in malaria parasites. Global response of selected peptides upon treatment with Cpd 9 (**A**) or Cpd 55 (**B**). Peptides with haemoglobin matching sequences as well as those which most likely derive from the same (24, 25) are highlighted with blue rectangle shapes. Fold changes relative to the DMSO control are means from 2 biological repeats collected in triplicate at each time of drug incubation.

### GNF179 elicits minimal discernible impact on the metabolome

KAF156 belongs to the imidazolopiperazine class of compounds that have been developed by the NITD and have shown potential as antimalarial agents for use in malaria treatment, prophylaxis and transmission blocking (34, 51). The exact mode of action of KAF156 is currently unknown but mutations in the *P. falciparum* cyclic amine resistance locus (PfCARL) as well as UDP-galactose and acetyl-CoA transporters have all been shown individually to confer resistance to KAF156 and its close analogues (33, 52). In our screen, GNF179 did not induce any significant metabolic effect after incubating parasites with the compound for 2.5 hours consistent with its slower rate of kill. Some low-level increase in purine metabolites (**Fig. 5A**) was observed along with a low-level accumulation of central carbon metabolism metabolites (malate, succinate and oxoglutarate) (**Fig. 5B**). This is also in agreement with previously reported metabolomics profiles of KAF156 as no significant changes in the parasite metabolome were observed after the same period of drug incubation (23). It is, therefore, not possible to predict the mode of action of GNF179 based on these profiles. Nevertheless, GNF179 is a relatively slow acting drug (**Fig. 1A**) which would suggest that extending exposure beyond 2.5 hours may be required to elicit an observable biochemical response, although ATQ (with a similar slow acting phenotype), induces a significant metabolic signal indicative of its mode of action over the same duration of drug exposure (2.5 hours). Future work using a longer incubation period might reveal unique signatures specific to this compound mode of action. Recently, *in vitro* selection for resistance and forward genetic screens have pointed to inhibition of protein trafficking as a possible mode of action of KAF156 and other imidazolopiperazines (53), so rapid/early metabolic changes might not be associated with mode of action.

**Fig. 5.**
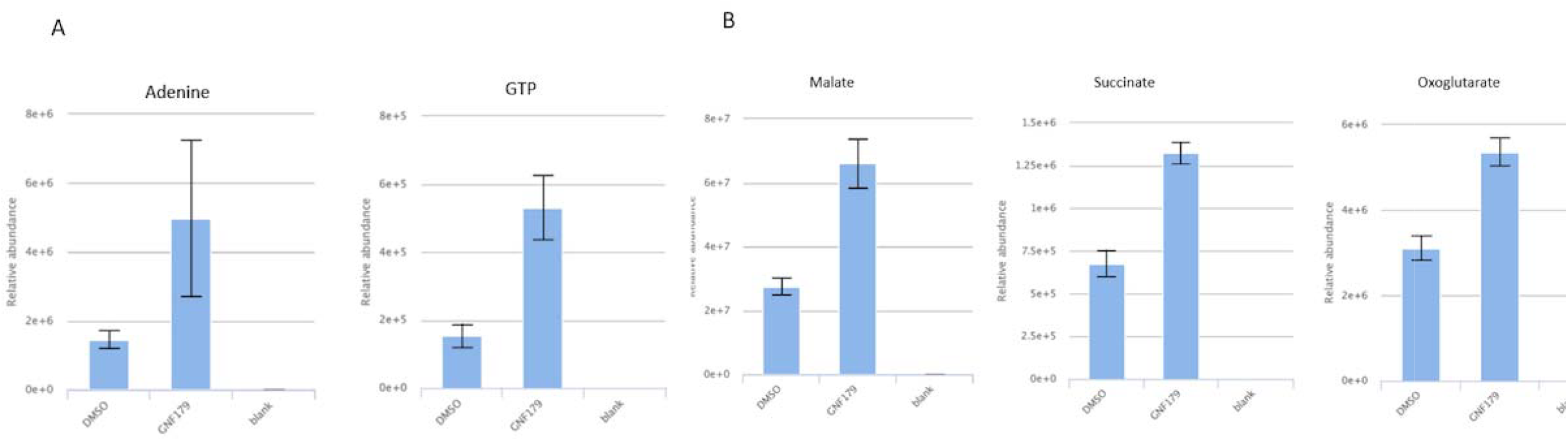
GNF179 metabolomic response for a selected metabolites. Relative abundance of adenine, GTP (**A**) and central carbon metabolism intermediates(**B**) in GNF179 treated parasites as compared to DMSO. Relative abundance comparisons of total ion counts are means from two biological repeats collected in triplicate at each time of drug incubation.

The initial global metabolomic responses to DHA, Cpd 55 and NITD246 are immediate, unique and dynamic Fast acting compounds may exert a parasite killing event in seconds or minutes with resulting metabolic profiles over time resulting from deregulated metabolic cascades which may or may not relate to a specific mode of action of the compounds. For example, the spiroindolone, KAE609 has been shown to lead to a rapid influx of sodium disrupting parasite ion homeostasis within seconds of drug exposure (44, 54). DHA is also known to target immediately several parasite proteins simultaneously in a promiscuous targeting process which leads to parasite death as a result of a disruption in several biological pathways (55, 56). In such events, metabolic and biochemical perturbation in essential pathways that are directly or indirectly involved in the mode of action of the compounds should be quantifiable within minutes of drug exposure. To this end, we aimed to resolve the dynamic metaprints of *P. falciparum* in response to the three fast-acting compounds studied here (DHA, Cpd 55 and NITD246) at 10 x IC_50_ exposure for 30 minutes, 1 hour and 2 hours. For all three compounds, significant changes to the global parasite metabolome were already observed after 30 minutes of drug exposure. Over the time course, the profile, extent and rate of change was time dependent and unique to each drug. The extent of change, however, was broadly equivalent for all three drugs after 2 hours. Of the ∼3000 mass features which were detected in each analysis by LC-MS, 4.3%, 1.6% and 4.6% changed significantly (>log2 fold change relative to DMSO control, adjusted p-value <0.05) at 30 minutes which increased to 5.6%, 5.8% and 5.2% at 1 hour and 7.8%, 8.1% and 7.9% at 2 hours for DHA, NITD246 and Cpd 55 respectively. Analysis of the volcano plots (**Fig. 6**) revealed that DHA elicits a stronger general downregulation of impacted metabolites after 30 minutes, although the magnitude of change converges towards those of NITD246 after 1 and 2 hours. The global metabolomic response to Cpd 55 revealed differences compared to DHA and NITD246 at all time points displaying lower magnitude fold changes symmetrically distributed around zero (**Fig. 6**). A unique, unvarying cluster of metabolites, possibly drug related adducts could be detected across the three time points (circled in red) in the Cpd 55 global metabolome. Overall, these global metabolome profiles indicate that, in spite of displaying a similar fast killing rate, and a signature impact on haemoglobin catabolism, additional discriminating features are already visible by 30 minutes of drug incubation and could point to as yet unknown drug-specific, parasite kill events.

**Fig. 6:**
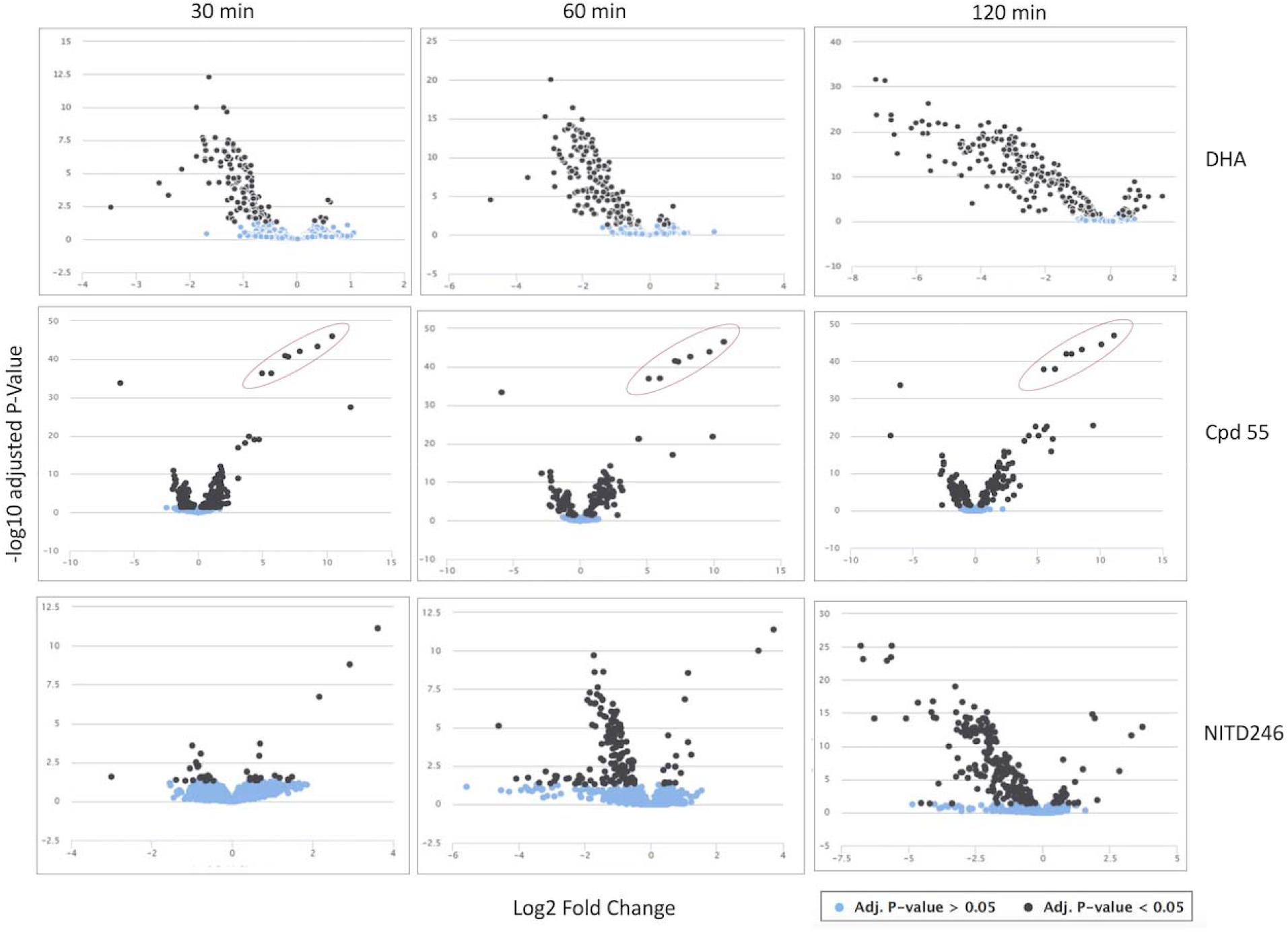
Global volcano plots of all detected mass features in DHA, Cpd 55 and NITD246 treatments according to their fold change relative to DMSO treatments. Significant features are represented as black dots while non-significant features are in light blue. n=2 with three technical replicates at each biological repeat.

### NITD246-induced perturbation of choline, pyrimidine and purine metabolites occurs shortly after compound administration

Our initial analyses established that NITD246 significantly perturbed metabolites in the pyrimidine, purine nucleotide and choline biosynthetic pathways after 2.5 hours of drug incubation (**Fig. 3, Fig. S2**). Analysis of the early time-resolved profiles of these metabolites permitted discrimination between a progressive and dynamic response or an immediate metabolic shock response. From these analyses, it was evident that NITD246 induced a gradual accumulation of choline related metabolites (**Fig. 7A**) that becomes visible at the 1- and 2-hour time points. Interestingly, DHA elicits a similar choline response profile (**Fig. S3A**) which may indicate a common consequence to primary target inhibition by both compounds. By contrast, Cpd 55 does not alter choline homeostasis (**Fig. 7A**), further suggesting that the metabolic consequences of DHA and NITD246 exposure in malaria parasites overlap while Cpd 55 acts via a different mode of action. A similar trend was observed for pyrimidine and purine nucleotide related metabolites that gradually decrease over 2 hours of NITD246 drug incubation but remain unchanged in Cpd 55 (**Fig. 7B-F**). Uniquely, orotate pools sharply accumulated in NITD246 treated parasites after 30 minutes of drug incubation but rapidly declined thereafter to significantly lower levels after 2 hours (**Fig. 7B**). This could be a result of preferential inhibition of the orotate phosphoribosyltransferase enzyme in the pyrimidine pathway during the initial assault on parasite ion homeostasis which would lead to accumulation of orotate that returns to normal levels as other enzymes of the pathway respond to the perturbation. DHA did not induce any significant alteration in either purine nucleotides or metabolites of the pyrimidine biosynthesis pathway during the 2-hour time period **(Fig. S3B-F**). Even though pyrimidine metabolite profiles were previously reported to change following exposure to DHA (10 x IC_50_, significant changes at 4 and 6 hours) (24), the observed absence of the pyrimidine profile with DHA in this study could be due to the shorter duration of drug exposure.

**Fig. 7.**
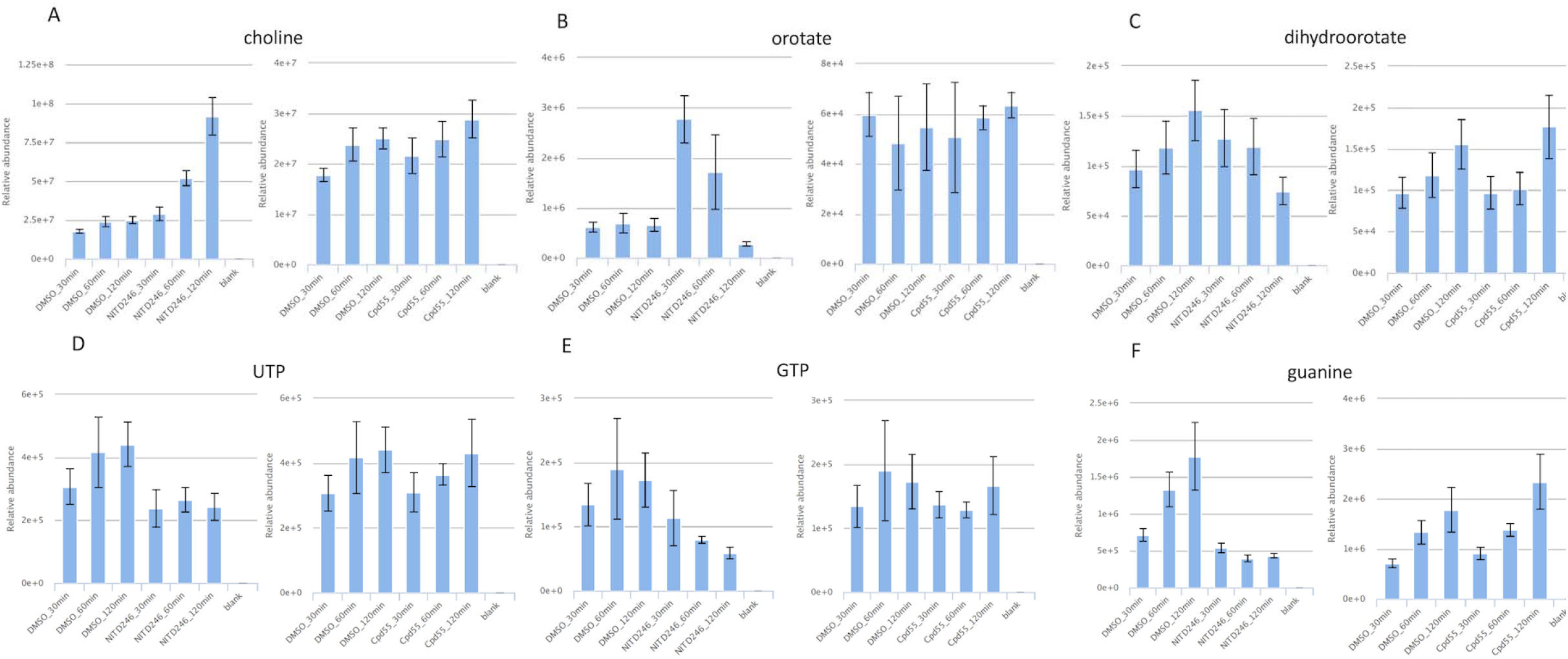
Time course comparisons of choline, pyrimidine and purine metabolites in DHA, NITD246 and Cpd 55 parasite treatments. Relative abundance of choline (**A**), selected indicated pyrimidine (**B-D**) and purine (**E, F**) metabolites. Comparisons of total ion counts are means from 2 biological repeats collected in triplicate at each time of drug incubation.

### Time dependent changes in peptide response for NITD246, DHA and Cpd 55

We next profiled the dynamic peptide response in NITD246, DHA and Cpd 55 treatments over the 0.5, 1 and 2 hours of drug incubation. Clustering of the global peptide response in the three drug treatments revealed that this response is already visible at 30 minutes of drug incubation and becomes more pronounced at 1 and 2 hours respectively (**Fig. 8A, Table S4**). Moreover, even though the global responses appear similar across the three compounds, Cpd 55 elicits a unique and early peptide response as compared to DHA and NITD246 which suggests a unique targeting of the parasite’s haemoglobin catabolism by this class of compound that does not directly or indirectly impact other biochemical pathways over the time-courses studied. Indeed, supervised clustering of the global peptidomes in these treatments by principal component analysis (PCA) revealed that at both time points, DHA and NITD246 peptide response clustered closer together than Cpd 55 or DMSO in the first and second principal components (**Fig. 8B**). Finally, some of the potentially haemoglobin derived peptides such Trp-Gly showed a time-dependent decline in NITD246 and Cpd 55 treatments (**Fig. 8C**) further suggestive of a global decline in parasite haemoglobin catabolism as a consequence of drug treatment.

**Fig 8:**
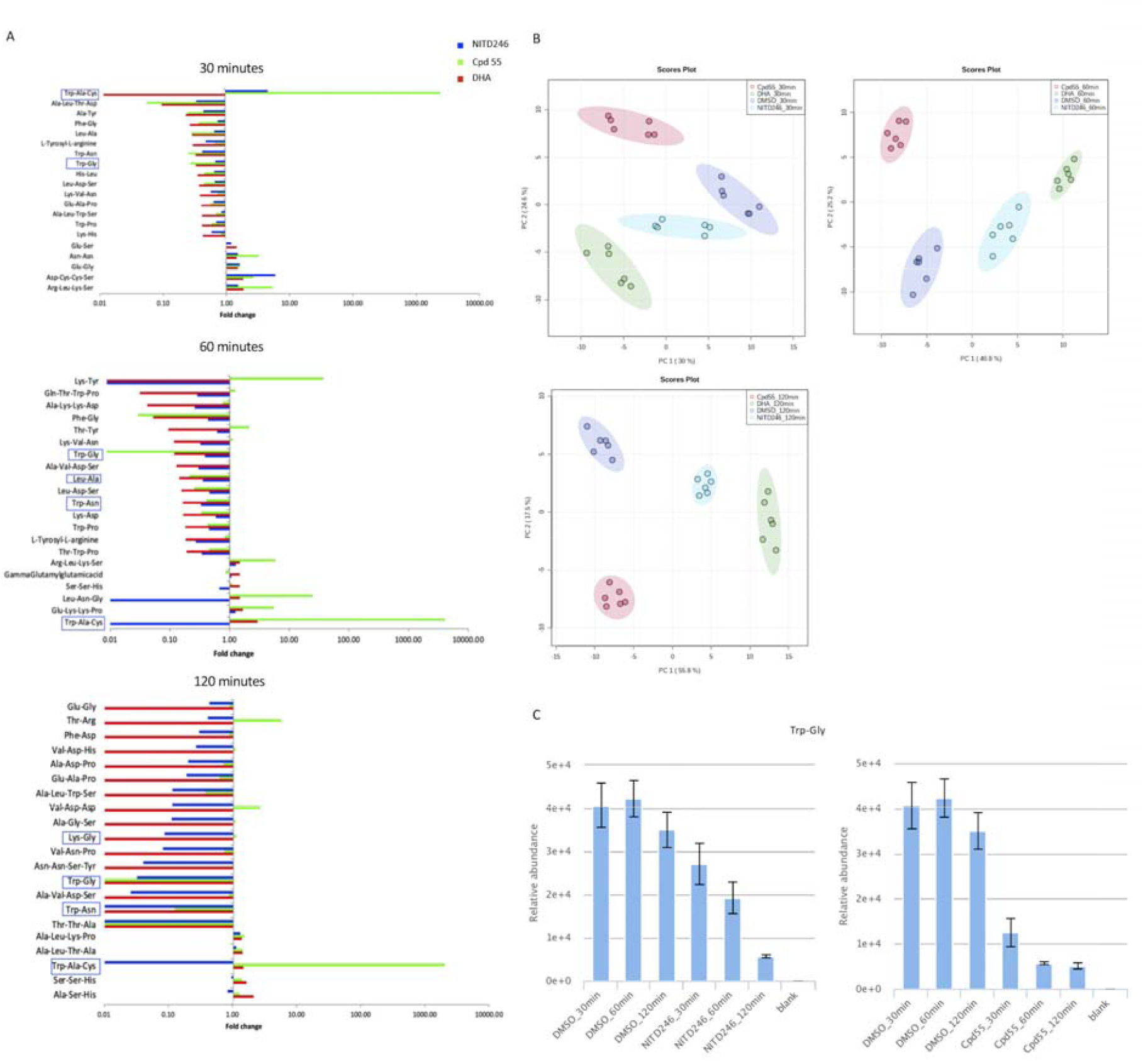
Time point resolution of global peptide responses in NITD246, DHA and Cpd 55 treated parasites. **A**. Global responses of top 20 significantly changed peptides upon treatment with DHA, NITD246 and Cpd 55 for 30 minutes, 1 and 2 hours. Peptides with haemoglobin matching sequences as well as those which have been previously validated to be haemoglobin derived (24, 25) are highlighted. **B**. PCA plots of the peptidomes of the three compounds after treatment for 30 minutes, 1 and 2 hours. **C** Relative abundance of a potential haemoglobin derived peptide Trp-Gly in DMSO vs the two indicated treatment compounds at the three time points. Fold changes and relative abundance comparisons are means from two biological repeats collected in triplicate at each time of drug incubation. PCA plots were carried out on log transformed data in Metaboanalysit 3 (66). 95% confidence intervals for each treatment group in the PCAs are highlighted with the indicated colors.

## Conclusions

In conclusion, this study reports the metabolomic profile of a novel class of fast acting compounds belonging to the ITD class which suggests haemoglobin catabolism as their possible mode of action or an essential event that precedes their parasite killing mechanism. By direct comparison with DHA and spiroindolones, we show that fast acting compounds elicit a unique, but broadly similar, peptide profile which could form a useful biochemical metaprint to identify fast acting compounds. Our approach to measure the time course of dynamic change of metabolome profiles shortly after drug administration may also help in differentiating fast acting compounds based on peptide clusters if the specific mechanism of action is unknown, in the context of selecting partner drugs for combinations to mitigate the emergence of resistance.

## Methods

### *P. falciparum* culture and maintenance

Two *P. falciparum* lines, 3D7 and a 3D7 derived luciferase reporter line (unpublished), were used for the experiments. The lines were cultured and maintained at 1-5% parasitaemia in fresh group O-positive red blood cells re-suspended to a 5% haematocrit in custom reconstituted RPMI 1640 complete media (Thermo Scientific) containing 0.23% sodium bicarbonate, 0.4% D-glucose, 0.005% hypoxanthine 0.6% Hepes, 0.5% Albumax II, 0.03% L-glutamine and 25mg/L gentamicin. Culture flasks were gassed with a mixture of 1% O_2_, 5% CO_2_, and 94% N_2_ and incubated at 37° C. For metabolomics experiments, parasites were scaled up to a 5-7% parasitaemia and sorbitol synchronized over 2 developmental cycles as previously described (57) to obtain parasites within a 3-6 hours developmental window. In brief, cultures were pelleted by centrifugation at 1,600 rpm for 3 min and resuspended in a 10× volume of 5% sorbitol (Sigma), followed by incubation at 37°C for 10 min. Following incubation, the infected RBCs cells were pelleted as above and washed in 40 pellet volumes of complete media before placing the infected RBCs back in fresh media and subsequent incubation at 37°C. Cultures were kept mycoplasma free and mycoplasma contaminations were monitored weekly using the Mycoalert Luminescence Kit (Lonza).

### *P. falciparum* SYBR Green I^®^ assay for parasite growth inhibition

Asynchronous stock cultures containing mainly ring stages were synchronised with 5% sorbitol as described above. Parasitaemia was quantified in the synchronized cultures with drug assays performed when the parasitaemia was between 1.5-5% with >90% rings. The stock culture was diluted to a haematocrit of 4% and 0.3% parasitaemia in complete media following which 50 μl was mixed with 50 μl of serial diluted drugs/inhibitors in complete media pre-dispensed in black 96 well optical culture plates (Thermo scientific) for a final haematocrit of 2%. Plates were gassed and incubated at 37° C for 72 hours followed by freezing at -20° C for at least 24 hours. The plate setup also included no drug controls as well as uninfected red cells at 2% haematocrit. After 72 hours of incubation and at least overnight freezing at -20° C, plates were thawed at room temperature for ∼4 hours. This was followed by addition of 100 μl to each well of 1X SYBR Green I^®^ (Invitrogen) lysis buffer containing 20 mM Tris, 5 mM EDTA, 0.008% saponin and 0.08% Triton X-100. Plate contents were mixed thoroughly by shaking at 700 rpm for 5 minutes and incubated for 1 hour at room temperature in the dark. After incubation, plates were read to quantify SYBR Green I^®^ fluorescence intensity in each well by a PHERAstar^®^ FSX microplate reader (BMG Labtech) or the CLARIOstar microplate reader (BMG Labtech) with excitation and emission wavelengths of 485 nm and 520 nm respectively. To determine growth inhibition, background fluorescence intensity from uninfected red cells was subtracted first. Fluorescence intensity of no drug controls was then set to correspond to 100% and subsequent intensity in presence of drug/inhibitor was calculated accordingly. Dose response curves and IC_50_ concentrations were plotted in Graph-pad Prism 7.

### *P. falciparum* luciferase assay

Metabolomics screens to characterise mode of action of compounds in malaria parasites rely on adequate exposure of parasites to the drugs for a good metabolic signal while avoiding over-exposure which can lead to death related metabolic signatures (23). This is however, difficult to quantify especially for fast acting compounds which could potentially elicit metabolic signatures very early on in the parasite killing cascade. Previous studies have employed a concurrent monitoring of glycolytic intermediates over the 6-hour course of drug exposure in mid-trophozoites as a marker of parasite viability (25). Nevertheless, we aimed to determine the killing kinetics of our compounds of interest biochemically by monitoring luciferase expression in a 3D7 dual reporter line that expresses NanoLuc and luciferase (3D7 luc, unpublished) under the control of a constitutive calmodulin promoter. Synchronised trophozoites (∼30 hours old) at 2% haematocrit and 2% parasitaemia were incubated with the compounds at 10X IC_50_ for 0.5,1,1.5, 2, 2.5, 3,3.5, 4, 5, 6 hours. Luciferase expression was quantified on a CLARIOstar microplate reader (BMG Labtech). Briefly, 100 ul of the reconstituted Dual-Luciferase^®^ Reporter reagent (Promega) was mixed with 100 ul parasite culture and incubated at room temperature in the dark for 15 minutes. Luciferase signal was quantified immediately after the incubation. Parasite viability was also monitored by microscopy analysis of methanol fixed Giemsa-stained smears.

### Magnetic purification of trophozoites

Drug induced metabolomics screens are mostly performed on 24–30-hour old trophozoites as they yield better metabolic signatures as well as less variability (23). To enrich for ∼24-30 hour old trophozoites, a magnetic separation was employed as previously described (58). Custom 3D printed magnet stands were assembled based on previously reported designs (58) and used to assemble a magnetic apparatus which was used to enrich for mature trophozoites in conjunction with cell separation LD columns (Miltenyi Biotech). Briefly, synchronized cultures at 5-7% parasitaemia (∼24-30 hours old) were re-suspended to 8% haematocrit following which 5 ml was loaded into the LD columns on the magnetic stands and allowed to flow through. Uninfected and early-stage parasites were washed off by loading the LD column with 5ml of clean complete media which allows for removal of all unbound RBCs. Bound parasites were then eluted in 5ml of fresh complete media after removal of the LD columns from the magnetic stands. Eluted parasites were pooled into a single falcon tube from which cell counts (haemocytometer counting) were performed and adjusted to a concentration of ∼1 × 10^8^ cells/ml. Purified parasites, containing >90% purified trophozoites were allowed to recover for ∼1 hour at 37° C at ∼0.5% haematocrit before the start of experiments. Further quality and purity of the enriched trophozoites was assessed by microscopy analysis of methanol fixed Giemsa-stained smears.

### Metabolite sample preparation

Magnetically purified trophozoites as described above were exposed to DHA, NITD246 and inactive analogue (NITD246i); GNF179; Cpd 9 and inactive analogue Cpd 9_ia; Cpd 55 and inactive analogue Cpd 55_ia at 10X IC_50_. ATQ was used as a positive control while DMSO (0.1%) was used in untreated controls. 1 ml of purified trophozoites (1 × 10 cells) was mixed with 4 ml of complete media containing spiroindolones, KAF156, ITDs and ATQ at 10 ^8^x IC_50_ in 6-well plates for 2.5 hours initially. The concentration used and the time of exposure was based on our time kill kinetics of these compounds as well as previously validated drug concentration and corresponding time points which are known to achieve a better metabolic signal resolution (23-25). To resolve the metabolic profiles of fast acting compounds at earlier time points, a similar approach as described above was used for DHA, NITD246 and Cpd 55 albeit with a dynamic drug exposure for 0.5, 1 and 2 hours. Incubations at all time points were performed in triplicate over 2 biological repeats. After drug incubation, 4 ml of media was aspirated from the 6-well plates and cells were resuspended in 1 ml volume and centrifuged to pellet the cells. Metabolism was immediately quenched by aspirating the supernatant and resuspending the cells in ice cold 1X PBS. All experiments were performed on ice onwards.

### Metabolite extraction

A mixture of water, methanol and chloroform (1:3:1) was used for metabolite extraction to allow for complimentary coverage of both polar and non-polar metabolites as previously described (59). The chilled suspension of cells was centrifuged at 8,500 g for 30 seconds at 4° C. After removing the supernatant, the cells were further washed by re-suspending in fresh 500 μl of ice cold 1X PBS and the supernatant was removed again. Cell pellets were then re-suspended in 200 μl of ice cold chloroform/methanol/water in a 1:3:1 ratio. After vigorously shaking for 1 hour in the cold room or chilled shaker at 4° C, the samples were sonicated for 2 minutes in ice-cold water and centrifuged at 15, 300g for 5 minutes at 4° C. ∼180ul of the supernatant was transferred to 2 ml clean screw capped tubes for LC-MS analysis. Pooled sample controls were also prepared during this time for quality control during the LC-MS processing. An extraction solvent blank was also included as part of the internal controls. Samples were kept at -80° C until processed.

### LC-MS Metabolomics analysis

Untargeted LC-MS sample processing was carried out at the University of Glasgow Polyomics facility on a hydrophilic interaction liquid chromatography (pHILIC) on a Dionex UltiMate 3000 RSLC system (Thermo Fisher Scientific) using a ZIC-pHILIC column (150 mm (length) × 4.6 mm (diameter), 5 μm (bead size) column) coupled to a Thermo Orbitrap Q-Exactive mass spectrometer (Thermo Fisher Scientific). 10 μl of the sample maintained on a 5° C auto-sampler was injected on a column that was maintained at 30 C. Samples were eluted on a linear gradient, starting with 20% A and 80% B for 15 min, followed by a 2 min wash with 95% A and 5% B, and 8 min re-equilibration with 20% A and 80% B, where solvent A is 20 mM ammonium carbonate in water while solvent B is acetonitrile. The LC-MS method was based on previously published protocols (60). Mass spectrometry was operated in polarity switching mode at a resolution of 70, 000, 10^6^ cts AGC target, spray voltages +⍰3.8 and −⍰3.8 kV, capillary temperature of 320 °C, heater temperature of 150 °C, sheath gas flow rate of 40 a.u., auxiliary gas flow rate of 5 a.u., sweep gas flow rate of 5 a.u., and a full scan mass window of 70–1050 m/z. m/z 83.0604, 149.0233 and 445.1200 were used as lock masses in the positive mode while m/z 89.0244 was used as a lock mass in the negative mode.

### Mass spectrometry fragmentation

Samples were also subjected to a fragmentation mass spectrometry analysis (LC-MS/MS) to allow for additional structural information on detected mass features. Fragmentation of the samples was carried out in either the positive or negative ionisation modes or both using duty cycles (1 full scan event and 1 top 5 or top 10 fragmentation event) as previously described (61).

### Data acquisition

Control runs consisting of blank runs and standardised internal controls were run in accordance with standard procedures at the Glasgow Polyomics to monitor the performance of the mass spectrometer in terms of chromatography and mass intensities. A mixture of standards containing 150 reference compounds available from Glasgow Polyomics were also run to assess the quality of the mass spectrometer and to aid in metabolite annotation and identification (60). Pooled samples containing fractional representation of samples were run prior to and across the batch every 6^th^ sample to monitor the stability and quality of the LC–MS run, whereas the actual samples were run in a randomised manner to minimise batch effects. Thermo Xcalibur Tune software was used for instrument control and data acquisition. After acquisition, all raw files were converted into mzXML format, separating positive and negative ionization mode spectra into two different mzXML files using the command line version of MSconvert (ProteoWizard).

### Data processing, analysis and metabolite identification

Data files in mzXML format were processed using an Excel interface, IDEOM (62), which is based on XCMS and mzmatch R tools that allow raw peak extraction, noise-filtering, gap-filling and peak annotations (63, 64). mzXML files were also processed using PiMP, a web based Glasgow Polyomics metabolomics data processing pipeline (65) that is also based on XCMS and mzmatch R tools (63, 64) but allows for easy and multiple sample comparisons across experimental conditions. Volcano plots and principal component analysis were visualised and plotted both in IDEOM, PiMP and Metaboanalyst 3 (66). Metabolite changes across different conditions and time points were plotted as fold changes or log2 fold changes. Identification of metabolites was based on fragmentation spectra, retention time and mass compared to authentic standards as previously outlined by the metabolomics standards initiative (MSI) (67). Metabolites that matched an authentic standard with or without a fragmentation spectrum were classified as identified (MSI level 1). Metabolites which did not match to any authentic standards but had spectral similarities with spectral libraries https://www.genome.jp/kegg/pathway.html, were classified as putatively annotated and analysed further based on fragmentation spectra if available.

## Acknowledgements

This work was supported by grants from the Wellcome Trust to A.P.W (083811/Z/07/Z; 107046/Z/15/Z). M.P.B. is funded by a Wellcome Trust core grant to the Wellcome Centre for Integrative Parasitology (104111/Z/14/Z). N.V.S was funded by a Commonwealth Doctoral Studentship (MWCS-2017-789) and the Novartis Global Health Fellowships.

