## Supplementary material for "A conserved metabolic signature associated with response to fast-acting antimalarial agents"

**Supplementary tables**

**Table S1: A global peptide response in KAE609a treated parasites as compared to DMSO at 2.5 hours** (separate excel sheet).

**Table S2: A global peptide response in Cpd 9 treated parasites as compared to DMSO at 2.5 hours** (separate excel sheet).

**Table S3: A global peptide response in Cpd 55 treated parasites as compared to DMSO at 2.5 hours** (separate excel sheet).

**Table S4: A global peptide response in KAE609a, Cpd 55 and DHA treated parasites as compared to DMSO at 30, 60 and 120 minutes** (separate excel sheet).

**List of supplementary figures**

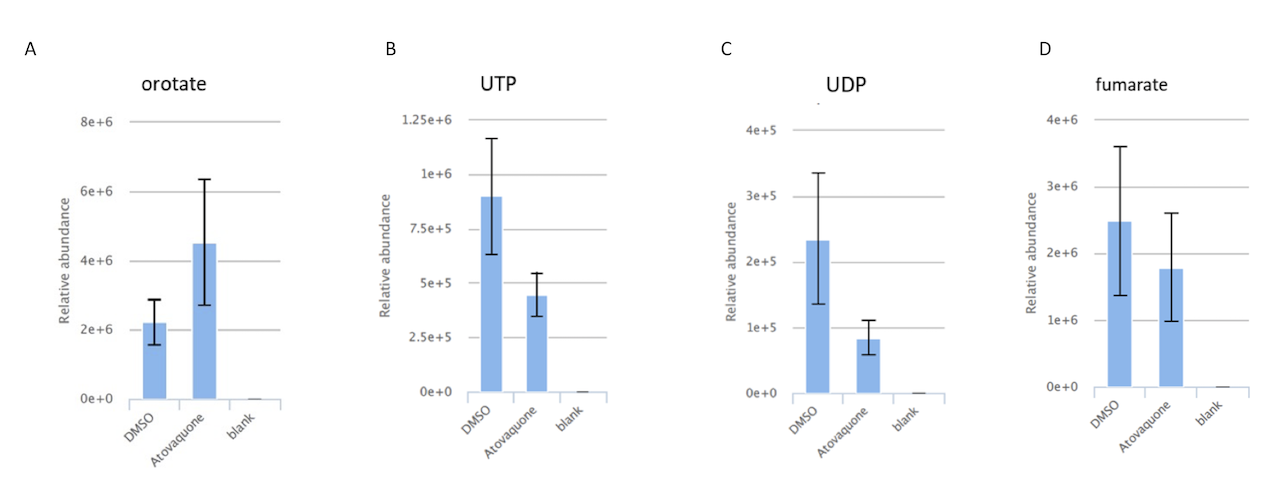

**Fig. S1. Pyrimidine and TCA cycle pathway metabolite responses to ATQ treatment.** Data was collected and processed as described in Fig. 2.

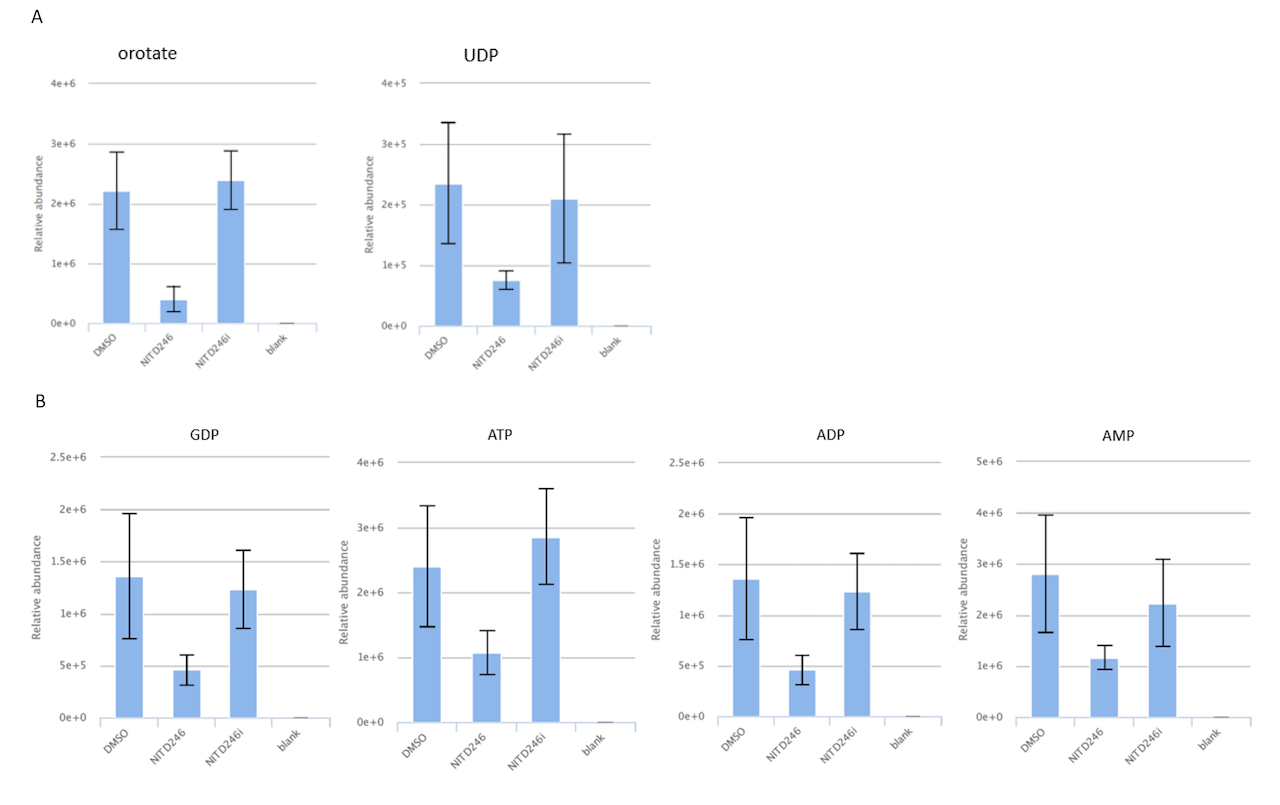

**Fig. S2**. **Relative abundance of indicated pyrimidine (A) and purine metabolites (B) metabolites in KAE609a treated parasites**. Data was collected and processed as described in Fig. 3.

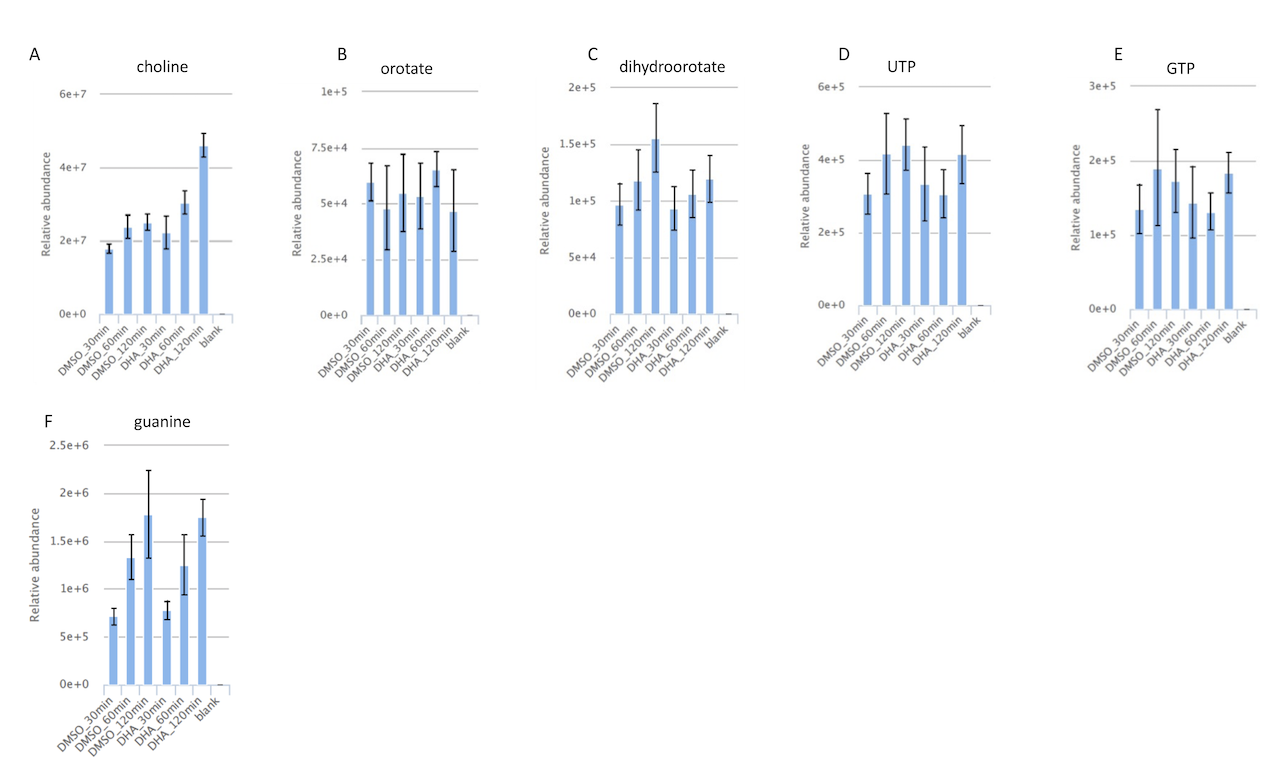

**Fig. S3. DHA choline, purine and pyrimidine responses in malaria parasites exposed to the drug for 30, 60 or 120 minutes.** Data was collected and processed as described in Fig. 7 for KAE609a and Cpd 55.
